## Supplementary Data for BioE3 for "BioE3 enables the identification of *bona fide* targets of E3 ligases"

Barroso-Gomila *et al*

SUPPLEMENTARY INFORMATION

**Supplementary Note 1:**

**Use of the wild type AviTag leads to general and nonspecific biotinylation**

To evaluate the spatial-specificity of the labelling obtained using the bioUb strategy, we first fused the $bio^{WHE}$ tag to a version of Ub that is not processable by DUBs (Ubnc for Ub non-cleavable, L73P mutation)[1]. This would avoid any recycling of locally pre-labelled $bio^{WHE}$Ubncs. Although this is not ideal and will disrupt regulation by DUBs, the bioUbnc is inducibly expressed for 24hrs only and likely does not reach levels of endogenous Ub. We generated HEK293FT and U2OS double stable cell lines for TRIPZ-$bio^{WHE}$Ubnc (puromycin-resistant) together with BirA alone or CEP120-BirA (blasticidin-resistant). The CEP120-BirA fusion protein localizes well to the centrosome. Doxycycline induction of $bio^{WHE}$Ubncs and 16 hours of biotin labelling yielded strong biotinylation of $bio^{WHE}$Ubncs in both conditions, with identical biotinylation patterns (Supplementary Fig. 1a). These results suggest that at longer biotin labelling times, BirA labels $bio^{WHE}$Ubncs in a general way, independently of their localization. To further evaluate the subcellular localization of the signal, we performed immunostainings on the U2OS double stable cell lines (with both inducible $bio^{WHE}$Ubnc and constitutive CEP120-BirA). The correct localization of CEP120-BirA was first confirmed, showing localization that correspond to centrosomes (single/paired dots, adjacent to or overlying the nucleus), while BirA alone localized to the nucleus and the cytoplasm (Supplementary Fig. 1b and c). We first observed that even without any biotin addition, CEP120-BirA and BirA carrying cell lines biotinylate $bio^{WHE}$Ubncs in a strong and general way when cultured with normal Fetal Bovine Serum (FBS) containing media (typically supplemented at 10%, which represents approximately 2 nM of biotin [2]; Supplementary Fig. 1b). However, when removing the biotin by dialyzing the serum, only the typical endogenous carboxylase biotinylation-derived streptavidin signal was observed (recognizable by the residual mitochondrial-like staining, Supplementary Fig. 1b). Strikingly, even when using dialyzed serum, biotin pulses as short as 2 minutes already showed a general labelling in the U2OS TRIPZ-$bio^{WHE}$Ubnc / BirA double stable cell line, whereas in the case of CEP120-BirA, the signal was localized at the centrosome (Supplementary Fig. 1c). Nevertheless, at longer timings, while CEP120-BirA remained specifically localized at the centrosome, we observed non-specific, general streptavidin localization after only 1 hour of biotin labelling (Supplementary Fig. 1c). Altogether, these results show that using dialyzed FBS enables the control of biotin labelling timings and that biotinylation of the $bio^{WHE}$ tag is unspecific to the localization of BirA enzyme even at short labelling timings, probably due to the high affinity between BirA and the bio$^{WHE}$ tag. In ohter words, non-conjugated bio$^{WHE}$UB is likely attracted to BirA and gets labelled, and incorporates into any substrates, primarily in the nucleus where abundant ubiquitination occurs. Further improvements were therefore required to achieve the specificity needed for BioE3 strategy. All the following experiments were performed using media supplemented with dialyzed (biotin-depleted) FBS.

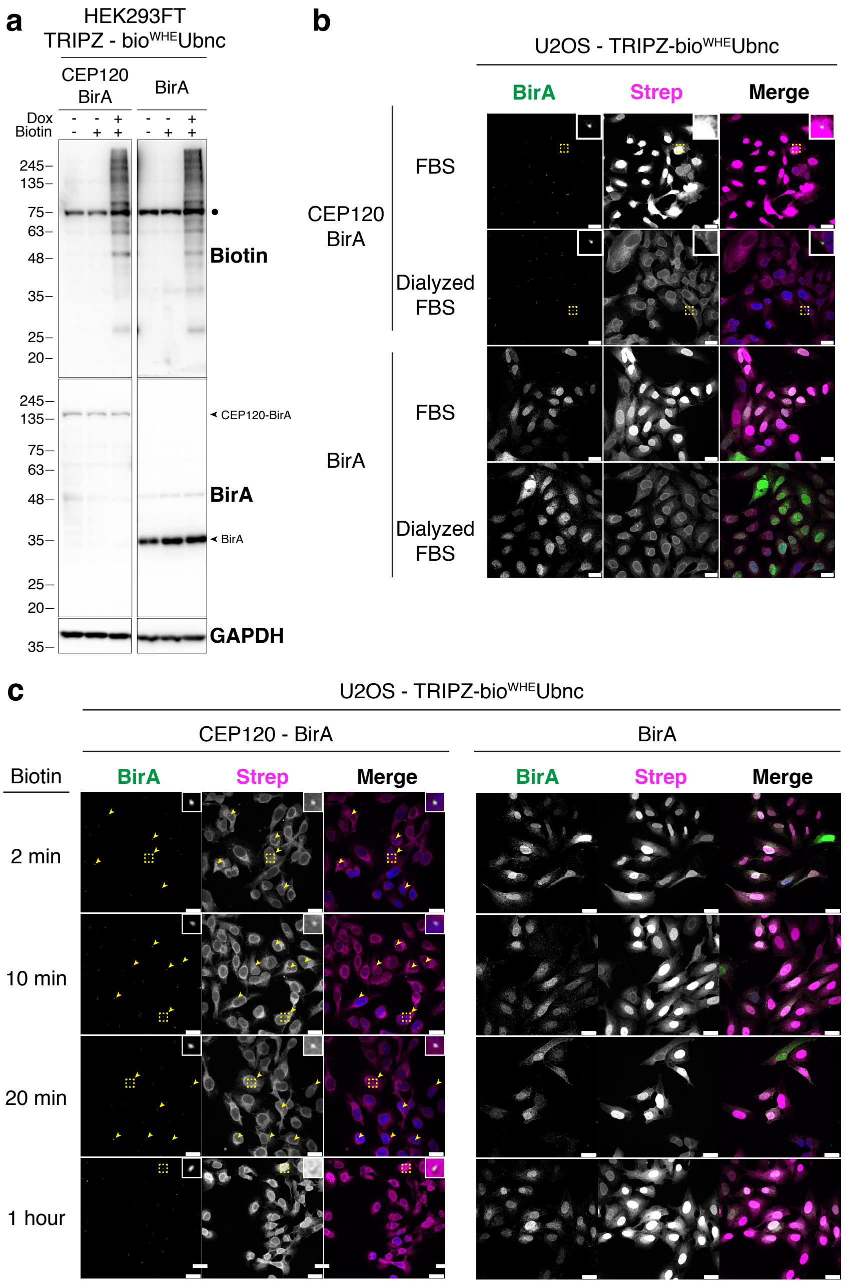


**<u>Supplementary Fig. 1:</u> Biotinylation of the wild type AviTag is general and unspecific. (a)** Western blot of 293FT stable cell lines expressing TRIPZ-bio$^{WHE}$Ubnc, together with EFS-BirA or the centrosome-localized CEP120-BirA. Doxycycline (DOX) induction was performed at 1 µg/ml for 24 hours and biotin supplementation at 50 µM for 16 hours. Similar general labelling of bio$^{WHE}$Ubncs was observed in both conditions. Dot indicates endogenous carboxylases that are biotinylated constitutively by the cell. Results are representative of three independent experiments performed on the same stable cell lines. Molecular weight markers are shown to the left of the blots in kDa. Source data are provided in the Source Data file. **(b-c)** Confocal microscopy of U2OS stable cell lines expressing TRIPZ-bio$^{WHE}$Ubnc, together with EFS-BirA or the centrosome-localized CEP120-BirA. Cells were pre-incubated in biotin-free dialyzed FBS-containing media or regular FBS-containing media for 24 hours prior to doxycycline induction at 1 µg/ml for 24 hours. No biotin was added in **(b)**. Cells cultured in dialyzed FBS only showed background originated from carboxylase biotinylation-derived streptavidin signal, while cells cultured in normal FBS showed general, unspecific labelling of bio$^{WHE}$Ubnc. 50 µM of biotin was added to dialyzed FBS-containing media cultured cells at indicated time-points in **(c)**. Colocalization of streptavidin and CEP120-BirA signals was observed at short biotin pulses (yellow arrowheads), while general unspecific labelling was observed at 1 hour of biotin treatment. Yellow dotted-line squares show the selected colocalization events for digital zooming. Nuclei are stained with DAPI (blue), biotinylated material with fluorescent streptavidin (Strep, magenta), and BirA with specific antibody (green). Black and white panels show the green and magenta channels individually. Results are representative of three independent experiments. Scale bar: 25 µm.

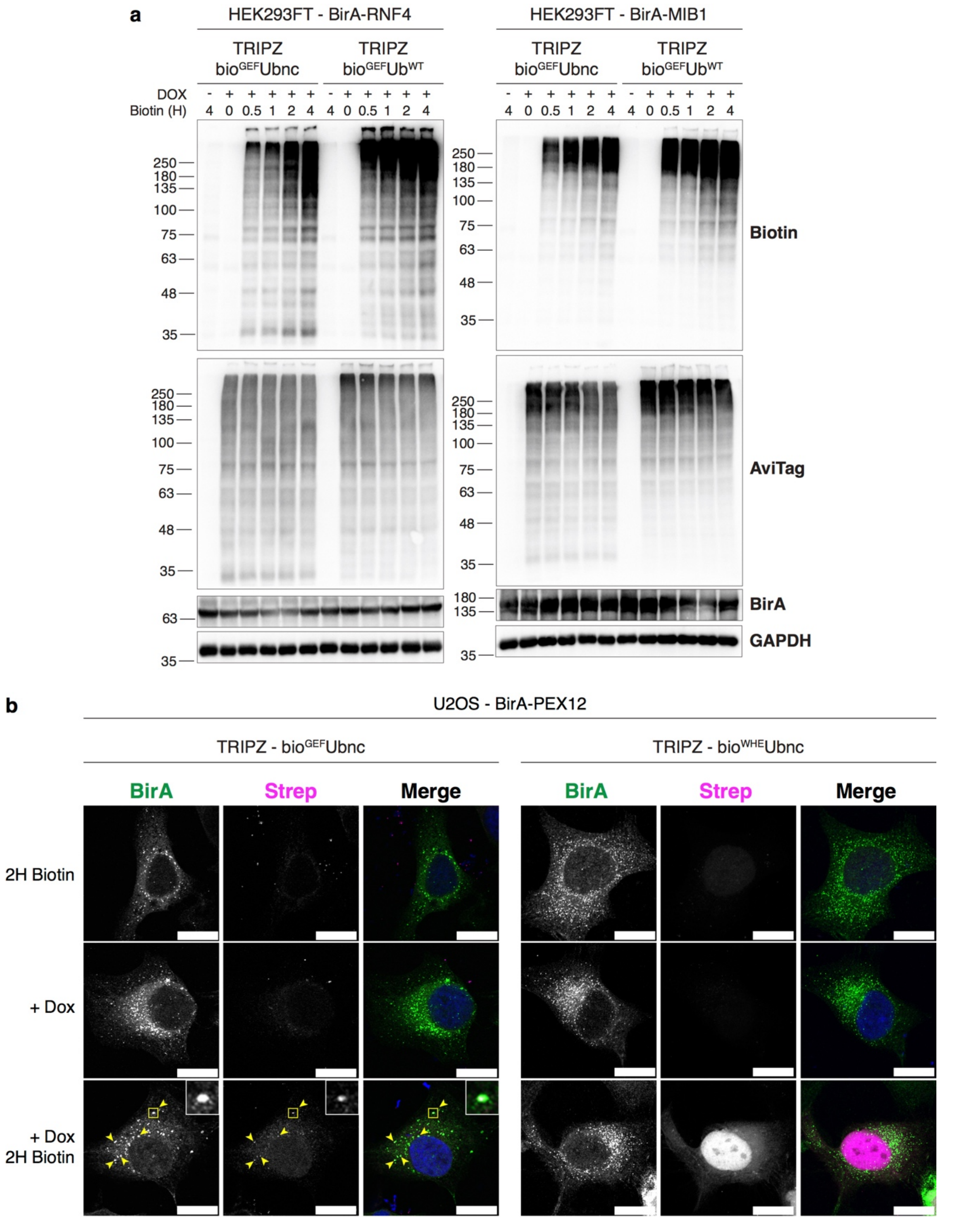


Supplementary Fig. 2: Low affinity bio$^{GEF}$ tag enables BioE3 studies. (a) Western blot of BioE3 experiment performed on HEK293FT stable cell lines expressing TRIPZ-bio$^{GEF}$Ubnc or TRIPZ-bio$^{GEF}$Ub$^{WT}$ and transfected with EFS-BirA-RNF4$^{WT}$ or EFS-BirA-MIB1$^{WT}$. Molecular weight markers are shown to the left of the blots in kDa. Cells were pre-incubated in dialyzed FBS-containing media prior to transfections, doxycycline (DOX) induction at 1 µg/ml for 24 hours and biotin supplementation at 50 µM for indicated time-points. Data are representative of 2 independent transfection experiments with similar results. Source data are provided in the Source Data file. (b) Confocal microscopy of U2OS stable cell lines expressing TRIPZ-bio$^{WHE}$Ubnc or TRIPZ-bio$^{GEF}$Ubnc transfected with EFS-BirA-PEX12. Cells were pre-incubated in biotin-free dialyzed FBS-containing media for 24 hours prior to transfection and doxycycline induction at 1 µg/ml for 24 hours. 50 µM of biotin was added for 2 hours. Correct colocalization of streptavidin and BirA-PEX12 signals was observed in peroxisomes for bio$^{GEF}$Ubnc (yellow arrowheads), while general unspecific labelling was detected for bio$^{WHE}$Ubnc. Dotted yellow line squares show the selected colocalization event for digital zooming. Nuclei are stained with DAPI (blue), biotinylated material with fluorescent streptavidin (Strep, magenta), and BirA with specific antibody (green). Black and white panels show the green and magenta channels individually. Scale bar: 10 µm.

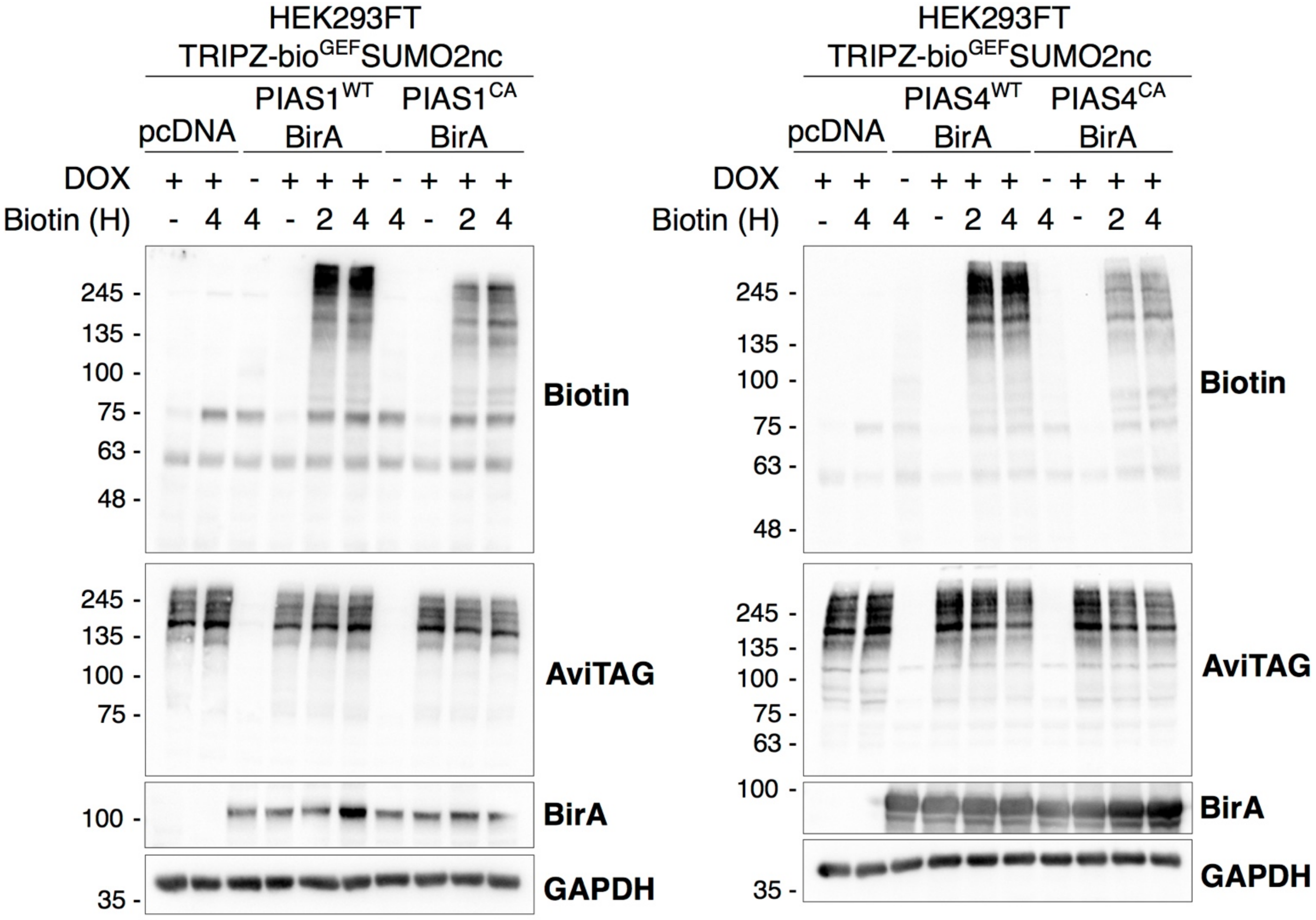


**Supplementary Fig. 3: BioE3 labels substrates of SUMO E3 ligases.** Western blot of BioE3 experiment performed on HEK293FT stable cell line expressing TRIPZ-bio$^{GEF}$SUMO2nc and transfected with CMV-PIAS1$^{WT}$-BirA or CMV-PIAS1$^{CA}$-BirA (left) and CMV-PIAS4$^{WT}$-BirA or CMV-PIAS4$^{CA}$-BirA (right). All BioE3 experiments were performed by pre-incubating the cells in dialyzed FBS-containing media prior to transfections, doxycycline (DOX) induction at 1 µg/ml for 24 hours and biotin supplementation at 50 µM for indicated time points. Molecular weight markers are shown to the left of the blots in kDa, antibodies used are indicated to the right. Data are representative of 2 independent transfection experiments with similar results. These samples were prepared for western analysis and not scaled-up for mass spectrometry analysis. Source data are provided in the Source Data file.

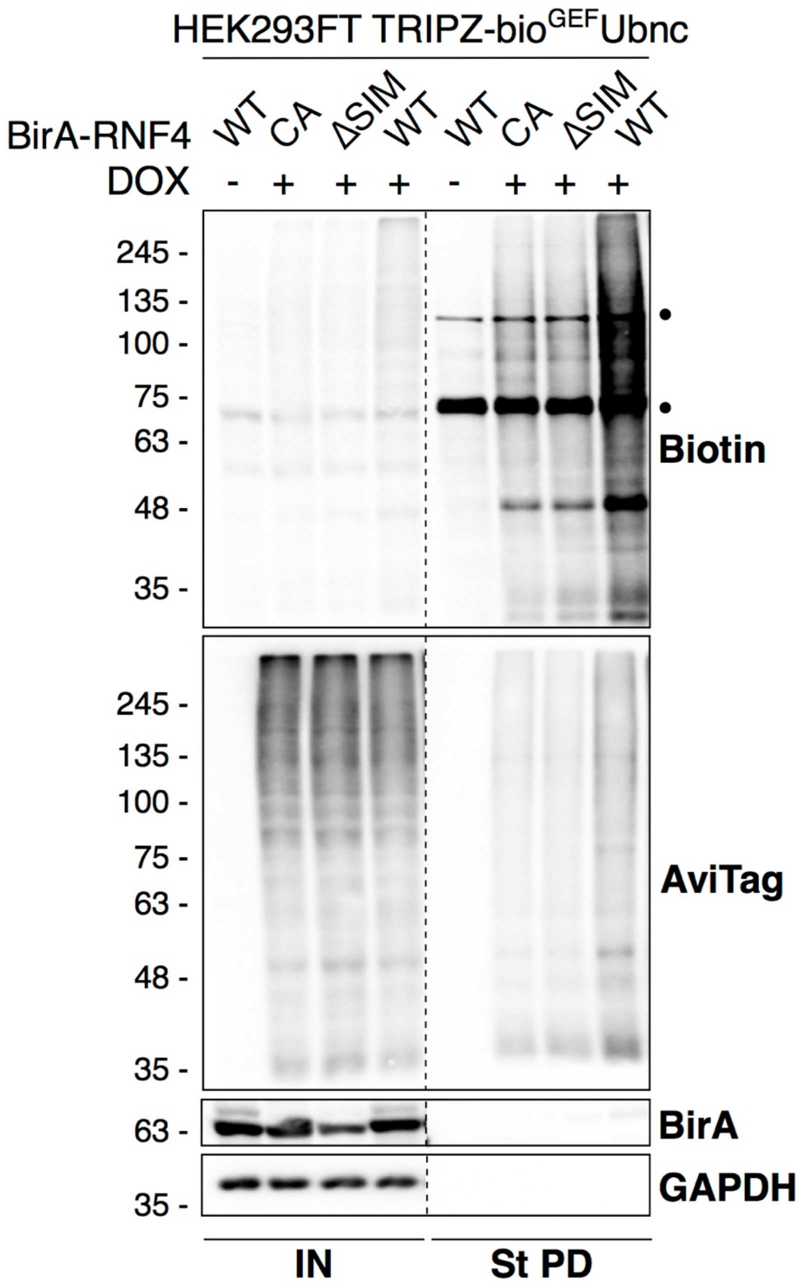


**<u>Supplementary Fig. 4:</u> BioE3 labels substrates of RNF4.** Western blot of BioE3 experiment performed on HEK293FT stable cell line expressing TRIPZ-bio$^{GEF}$Ubnc and transfected with EFS-BirA-RNF4$^{WT}$, BirA-RNF4$^{CA}$ or BirA-RNF4$^{\Delta SIM}$ (related to Fig. 4). IN: input; St PD: streptavidin pull-down. Dotted line indicates a cut in the same blot. Dots indicate endogenously biotinylated carboxylases. Molecular weight markers are shown to the left of the blots in kDa, antibodies used are indicated to the right. Data are representative of 3 independent transfection experiments with similar results. These samples are from scaled-up experiments for mass spectrometry analysis, and representative of the three ndependent replicates. Source data are provided in the Source Data file.

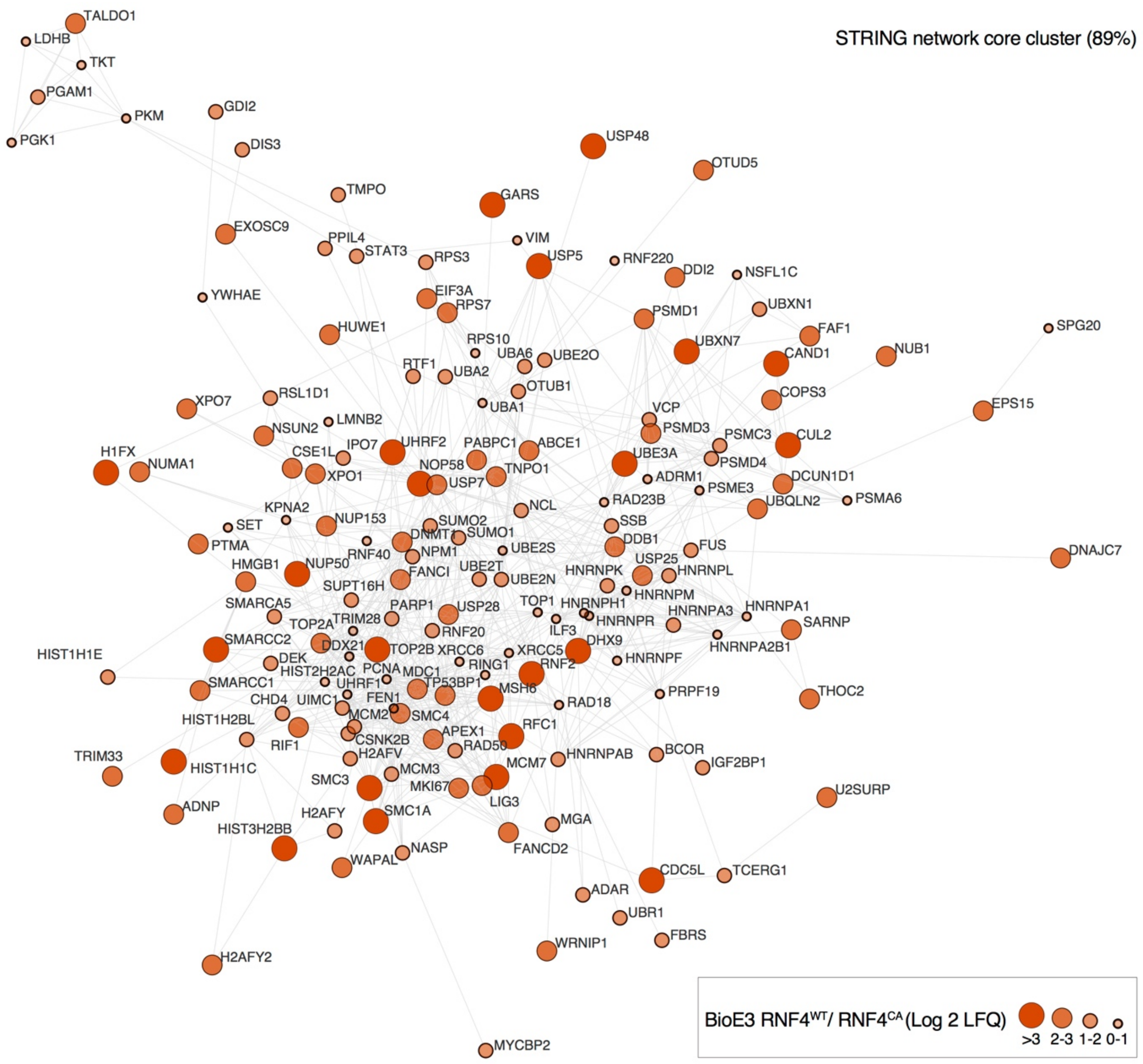


**<u>Supplementary Fig. 5:</u> RNF4 Ub substrates form a high interconnected core-cluster.** STRING network analysis of bio$^{GEF}$Ubnc RNF4 targets defined in Fig. 4a, showing a high interconnected network composed of the 89% of the proteins. Color, transparency and size of the nodes were discretely mapped to the Log2 enrichment value as described.

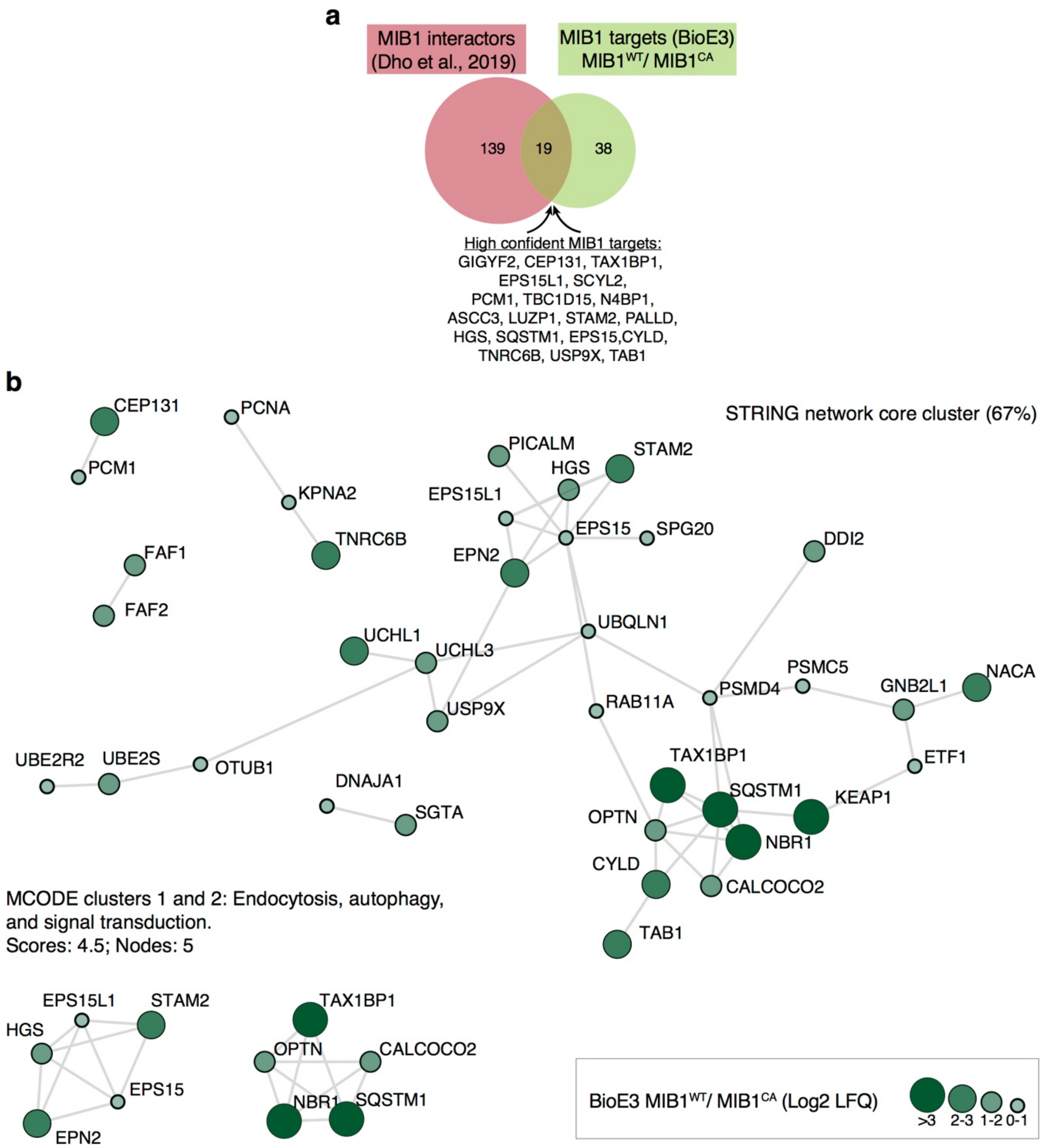


**<u>Supplementary Fig. 6:</u> MIB1 Ub targets participate in centrosomal proteostasis, Ub, proteasome, endocytosis and autophagy related processes. (a)** Venn diagram showing the Ub targets of MIB1 (comparison of the BioE3 MIB1$^{WT}$/MIB1$^{CA}$ targets in Fig. 6c) and the MIB1 interactome (MIB1 BioID from Dho *et al.* [3]). Comparisons data are provided in Supplementary Data 3. **(b)** STRING network analysis of bio$^{GEF}$Ubnc MIB1 targets defined in Fig. 6c, showing a high interconnected network composed of the 71% of the proteins. Highly interconnected sub-clusters were derived and characterized using MCODE. Color, transparency and size of the nodes were discretely mapped to the Log2 enrichment value as described.

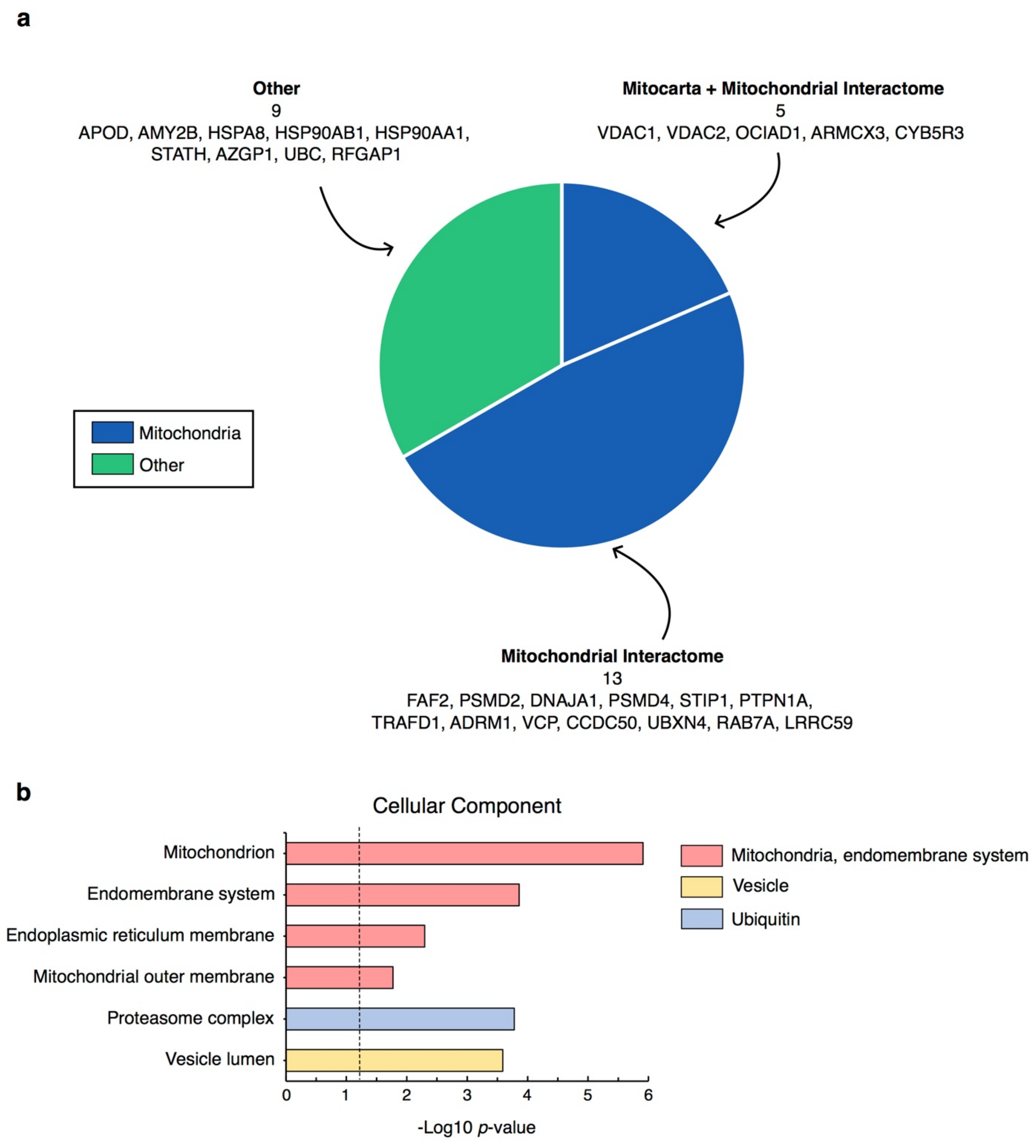


**Supplementary Fig. 7: MARCH5 BioE3 identifies mitochondrial proteins. (a)** Comparison of the BioE3 MARCH5 targets identified in Fig. 7e with Mitocarta [4], an inventory of mitochondrial proteins, and the mitochondrial proximity interaction network defined by Antonicka *et al.* [5]. Comparisons data are provided in Supplementary Data 5. **(b)** Gene ontology analysis of the MARCH5 targets defined in Fig. 7e. Depicted cellular components were significantly enriched. Dotted line represents the threshold of the *p*-value (0.05). Data are provided as Supplementary Data 6.

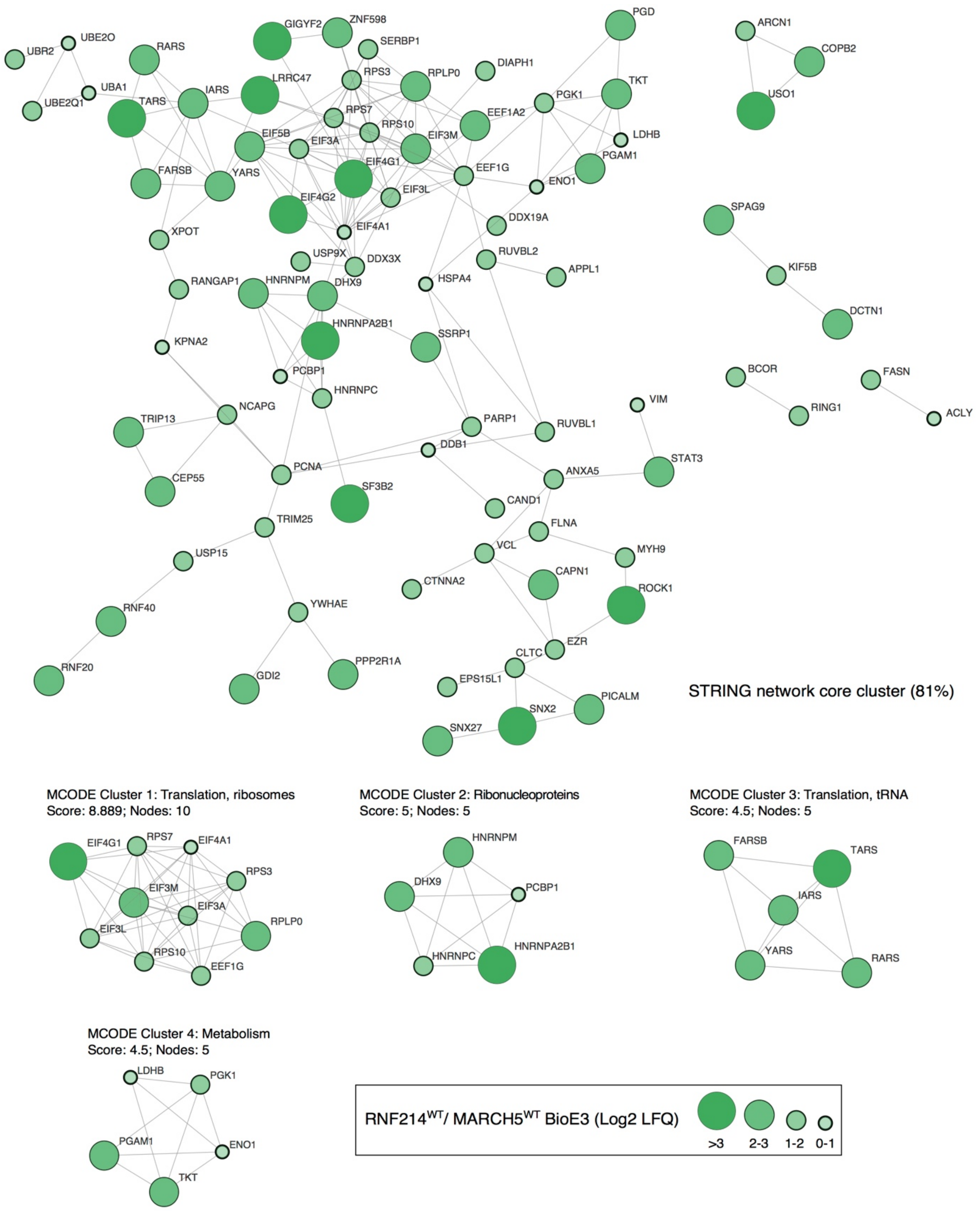


**<u>Supplementary Fig. 8:</u> RNF214 targets are involved in translation, intracellular trafficking and cytoskeleton.** STRING network analysis of the RNF214 targets defined in Fig. 7e, showing a high interconnected network composed of the 81% of the proteins. Highly interconnected sub-clusters were derived from the core-cluster using MCODE. Color, transparency and size of the nodes were discretely mapped to the Log2 enrichment value as described.

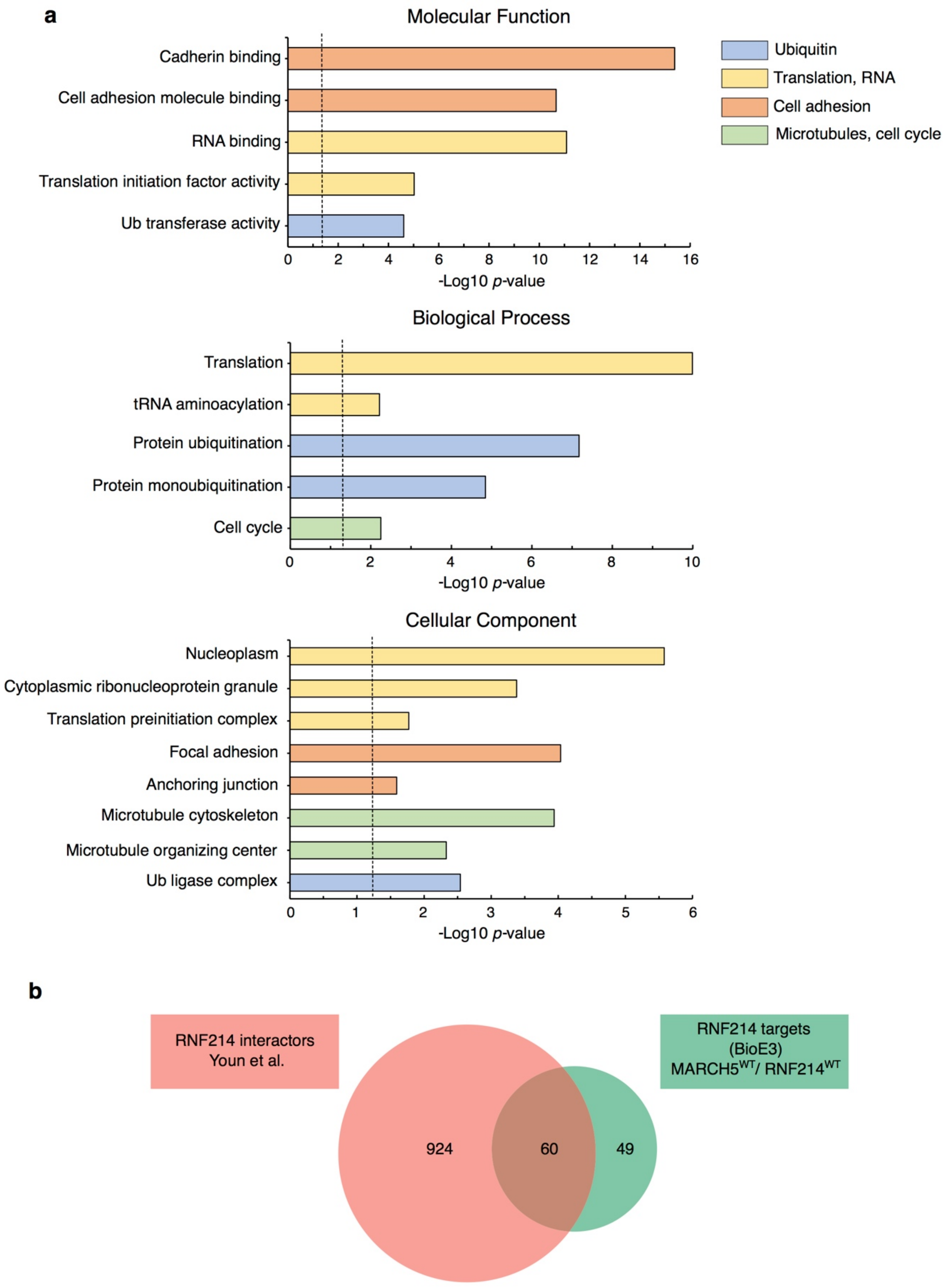


<u>Supplementary Fig. 9:</u> **BioE3 identifies RNF214 targets related to translation, actin cytoskeleton, microtubules and Ub. (a)** Gene ontology analysis of the RNF214 targets defined in Fig. 7e. Depicted biological processes, molecular functions and cellular components were significantly enriched. Dotted line represents the threshold of the *p*-value (0.05). Data are provided as Supplementary Data 6. **(b)** Venn diagram showing the Ub targets of RNF214 (comparison of the BioE3 MARCH5$^{WT}$/RNF214$^{WT}$ targets in Fig. 7e) and the RNF214 interactome (RNF214 BioID from Youn *et al.* [6]). Comparisons data are provided in Supplementary Data 5.

**Supplementary Note 2:**

**NEDD4 E3 ligase activation enhances BioE3 activity**

We evaluated NEDD4 BioE3 efficiency by performing standard BioE3 experiments comparing BirA-NEDD4$^{WT}$ and its transthiolation deficient BirA-NEDD4$^{CA}$ mutant. We observed very low and comparable levels of BioE3 biotinylation activity when using either BirA-NEDD4$^{WT}$ or BirA-NEDD4$^{CA}$ (Supplementary Fig. 10a, Biotin blot). This lack of BioE3 activity is probably due to the fact that NEDD4 ligases need to be activated, through EGF/FGF or intracellular calcium uptake, to stimulate their E3 ligase activity [7-9]. We thus evaluated NEDD4 Ub BioE3 activity in the U2OS – TRIPZ-bio$^{GEF}$Ubnc cell line, treating the cells with ionomycin and $CaCl_2$ to induce intracellular calcium uptake and with MG132 to inhibit proteasomal degradation. We observed that, at basal-conditions, inactivated NEDD4 localizes to the cytoplasm as reported previously, showing some BioE3 activity (Supplementary Fig. 10b). However, upon NEDD4 activation through ionomycin treatment, we observed that NEDD4 localized to the plasma membrane as well as to cytoplasmic structures that might correspond to vesicles, with high BioE3 activity that was accumulated upon proteasomal inhibition (Supplementary Fig. 10b). Thus, NEDD4 ligase needs to be activated to perform efficient BioE3 experiments. We therefore decided to mimic this activation process by mutating its autoinhibitory C2 domain. We generated a BirA-NEDD4$^{\Delta C2}$ version that lacks the entire C2 domain, as well as a BirA-NEDD4$^{3M}$ version in which three key amino acids that participate in NEDD4-closed conformation were mutated (I36A, L37A and Y604A) [10]. We evaluated NEDD4 Ub BioE3 using BirA- NEDD4$^{WT}$, NEDD4$^{CA}$, NEDD4$^{3M}$ or NEDD4$^{\Delta C2}$ and observed that the hyper-activated versions NEDD4$^{3M}$ and NEDD4$^{\Delta C2}$ showed enhanced BioE3 activity (Supplementary Fig. 10c). These results show that activation of NEDD4 is essential to detect its E3 ligase activity.

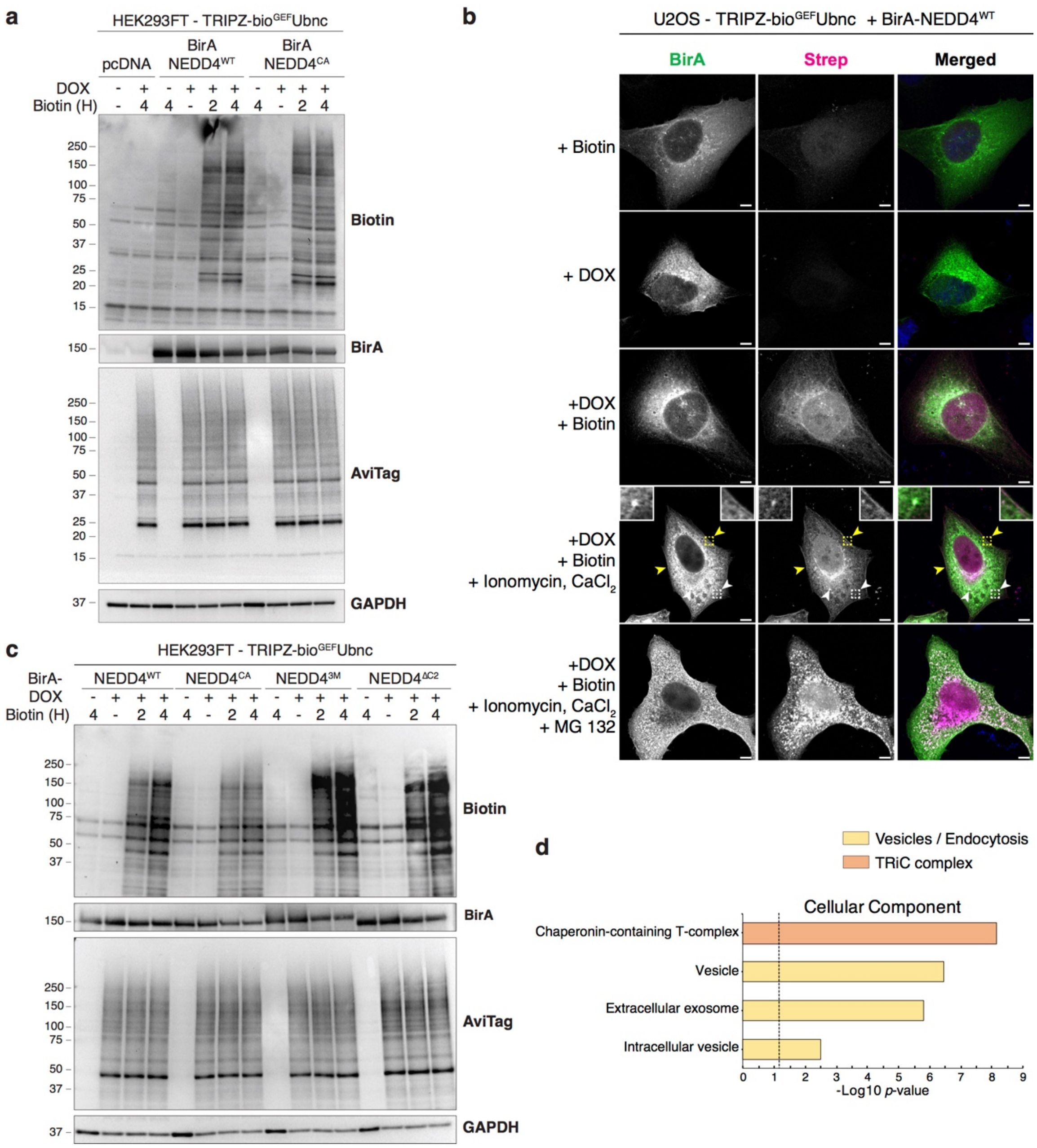


**Supplementary Fig. 10: NEDD4 E3 ligase activation enhances BioE3 activity. (a)** Western blot of BioE3 experiment performed on 293FT stable cell line expressing TRIPZ-bio$^{GEF}$Ubnc and transfected with EFS-BirA-NEDD4$^{WT}$ or BirA-NEDD4$^{CA}$. No efficient BioE3 activity was detected. **(b)** Confocal microscopy of BioE3 experiment performed on U2OS stable cell line expressing TRIPZ-bio$^{GEF}$Ubnc and transiently transfected with EFS-BirA-NEDD4$^{WT}$. Cells were also treated with 1 µM ionomycin and 2 mM $CaCl_2$ for 2 hours to induce intracellular calcium uptake and with 10 µM MG132 for 6 hours to inhibit proteasomal degradation. Colocalization of streptavidin and BirA-NEDD4$^{WT}$ was observed at plasma membrane (yellow arrowheads; dotted yellow line squares digital zooming, top right box) and intracellular vesicles (white arrowheads; dotted white line squares digital zooming, top left box) upon calcium-mediated activation of NEDD4$^{WT}$, and the signal accumulates upon MG132 treatment. Nuclei are stained with DAPI (blue), biotinylated material with fluorescent streptavidin (Strep, magenta), and BirA with specific antibody (green). Black and white panels show the green and magenta channels individually. Scale bar: 8 µm. **(c)** Western blot of BioE3 experiments performed on 293FT stable cell line expressing TRIPZ-bio$^{GEF}$Ubnc and transiently transfected with EFS-BirA-NEDD4$^{WT}$, BirA-NEDD4$^{CA}$, BirA-NEDD4$^{3M}$ or BirA-NEDD4$^{\Delta C2}$. NEDD4 activating mutations showed enhanced BioE3 activity. **(d)** Gene ontology analysis of the NEDD4 targets defined in Fig. 8g. Depicted cellular components were significantly enriched. Dotted line represents the threshold of the *p*-value (0.05). Data are provided as Supplementary Data 8. **(a-c)** Results are representative of two independent transfection experiments. Molecular weight markers are shown to the left of the blots in kDa. Source data are provided in the Source Data file.

**<u>Supplementary Table 1:</u> List of the constructs used in this study.** Related to Cloning section of *Methods*. The plasmid name, the resistance for bacterial transformation and amplification (Res), the plasmid backbone in which the DNA of interest (Plasmid insert) was inserted, and notes that explain the backbone as well as the source of the inserted DNA are depicted. The rows of the plasmids that are used afterwards as backbones are depicted in grey.

| Plasmid Name | Res | Plasmid backbone | Plasmid insert | Sources/cloning notes |
|---|---|---|---|---|
| EYFP-N1 | KAN | EYFP-N1 | | CMV- EYFP (EcoR1-Not1) |
| CMV-bio$^{WHE}$SUMO1 | KAN | EYFP-N1 | bio$^{WHE}$SUMO1$^{WT}$ (EcoR1-Not1) | Replaced EYFP (EcoR1-Not1) with AviTag-tagged SUMO1; SUMO1 source: hTERT-RPE1 cDNA |
| CMV-bio$^{WHE}$SUMO2 | KAN | EYFP-N1 | bio$^{WHE}$SUMO2$^{WT}$ (EcoR1-Not1) | Replaced EYFP (EcoR1-Not1) with AviTag-tagged SUMO2; SUMO2 source: hTERT-RPE1 cDNA |
| CMV-bio$^{WHE}$Ub | KAN | EYFP-N1 | bio$^{WHE}$Ub$^{WT}$ (EcoR1-Not1) | Replaced EYFP (EcoR1-Not1) with Avi-tagged ubiquitin (Ub); Ub source: CAG-bioUB [11] |
| CMV-bio$^{WHE}$SUMO1nc | KAN | EYFP-N1 | bio$^{WHE}$SUMO1nc (EcoR1-Not1) | Incorporates mutation Q94P to suppress deSUMOylation from substrates and terminates in GG to bypass initial protease activation step. Source: CMV-bio$^{WHE}$SUMO1. |
| CMV-bio$^{WHE}$SUMO2nc | KAN | EYFP-N1 | bio$^{WHE}$SUMO2nc (EcoR1-Not1) | Incorporates mutation Q90P to suppress deSUMOylation from substrates and terminates in GG to bypass initial protease activation step. Source: CMV-bio$^{WHE}$SUMO2. |
| CMV-bio$^{WHE}$Ubnc | KAN | EYFP-N1 | bio$^{WHE}$Ubnc (EcoR1-Not1) | Incorporates mutation L73P to suppress deubiquitylation from substrates and terminates in G-G to bypass initial protease activation step. Source: CMV-bio$^{WHE}$Ub. |
| CMV-bio$^{GEF}$SUMO1nc | KAN | EYFP-N1 | bio$^{GEF}$SUMO1nc (EcoR1-Not1) | Changed WHE to GEF in AviTag. Source: CMV-bio$^{WHE}$SUMO1nc. |
| CMV-bio$^{GEF}$SUMO2nc | KAN | EYFP-N1 | bio$^{GEF}$SUMO2nc (EcoR1-Not1) | Changed WHE to GEF in AviTag. Source: CMV-bio$^{WHE}$SUMO2nc |
| CMV-bio$^{GEF}$Ubnc | KAN | EYFP-N1 | bio$^{GEF}$Ubnc (EcoR1-Not1) | Changed WHE to GEF in AviTag. Source: CMV-bio$^{WHE}$Ubnc. |
| TRIPZ | AMP | TRIPZ-FF3 | TurboRFP and shRNAmir for firefly luciferase | Based on TRIPZ (lentiviral all-in-one doxycyline-inducible vector; OpenBiosystems/Thermo); Gibson-cloned elements into BshT1-Mlu1 digested vector (removing TurboRFP and shRNA). |
| TRIPZ-bio$^{WHE}$SUMO1nc-PURO | AMP | TRIPZ | bio$^{WHE}$SUMO1nc | Source: CMV-bio$^{WHE}$SUMO1nc. |
| TRIPZ-bio$^{WHE}$SUMO2nc-PURO | AMP | TRIPZ | bio$^{WHE}$SUMO2nc | Source: CMV-bio$^{WHE}$SUMO2nc |
| TRIPZ-bio$^{WHE}$Ubnc-PURO | AMP | TRIPZ | bio$^{WHE}$Ubnc | Source: CMV-bio$^{WHE}$Ubnc |
| TRIPZ-bio$^{GEF}$SUMO1nc-PURO | AMP | TRIPZ | bio$^{GEF}$SUMO1nc | Source: CMV-bio$^{GEF}$SUMO1nc |
| TRIPZ-bio$^{GEF}$SUMO2nc-PURO | AMP | TRIPZ | bio$^{GEF}$SUMO2nc | Source: CMV-bio$^{GEF}$SUMO2nc |
| TRIPZ-bio$^{GEF}$Ubnc-PURO | AMP | TRIPZ | bio$^{GEF}$Ubnc | Source: CMV-bio$^{GEF}$Ubnc |
| Lenti-EFS-GSQ-BirA-P2A-BLAST | AMP | Lenti-EFS-P2A-blast | BirA$^{WT/OPT}$ (EcoR1-Not1) | Based on Lenti-Cas9-blast; Cas9 removed BshT1-BamH1. Elements: EFS-BshT1-ASC1-GSQ-EcoR1-BirA$^{WT/OPT}$-Not1-BamH1-P2A-blast; BirA Source: pOZFHHN-HumBirA (gift of V. Ogryzko) |
| Lenti-EFS-CEP120-GS-BirA-P2A-BLAST | AMP | Lenti-EFS-GSQ-P2A-BLAST | CEP120-BirA (BshT1-EcoR1) | CEP120 Source: hTERT-RPE1 cDNA |
| Lenti-EFS-BirA-GSQ-PEX12-P2A-BLAST | AMP | Lenti-EFS-GSQ-BirA-P2A-BLAST | PEX12$^{WT}$ (EcoR1-Not1) | PEX12 Source: hTERT-RPE1 cDNA |
| Lenti-EFS-BirA-GSQ-PIAS1-P2A-BLAST | AMP | Lenti-EFS-GSQ-BirA-P2A-BLAST | PIAS1$^{WT}$ (EcoR1-Not1) | PIAS1 Source: hTERT-RPE1 cDNA |
| Lenti-EFS-BirA-GSQ-PIAS1$^{CA}$ -P2A-BLAST | AMP | Lenti-EFS-GSQ-BirA-P2A-BLAST | PIAS1$^{CA}$ (EcoR1-Not1) | Catalytically inactive: C351A mutation of RING domain. PIAS1 Source: hTERT-RPE1 cDNA |
| Lenti-EFS-BirA-GSQ-PIAS4-P2A-BLAST | AMP | Lenti-EFS-GSQ-BirA-P2A-BLAST | PIAS4$^{WT}$ (EcoR1-Not1) | PIAS4 Source: hTERT-RPE1 cDNA |
| Lenti-EFS-BirA-GSQ-PIAS4$^{CA}$ -P2A-BLAST | AMP | Lenti-EFS-GSQ-BirA-P2A-BLAST | PIAS4$^{CA}$ (EcoR1-Not1) | Catalytically inactive: C342A mutation of RING domain. PIAS4 Source: hTERT-RPE1 cDNA |
| Lenti-EFS-BirA-GSQ-RNF4-P2A-BLAST | AMP | Lenti-EFS-GSQ-BirA-P2A-BLAST | RNF4$^{WT}$ (BshT1-Asc1) | RNF4 Source: hTERT-RPE1 cDNA |
| Lenti-EFS-BirA-GSQ-RNF4$^{CA}$- P2A-BLAST | AMP | Lenti-EFS-GSQ-BirA-P2A-BLAST | RNF4$^{CA}$ (BshT1-Asc1) | Catalytically inactive: C159A mutation of RING domain. RNF4 Source: hTERT-RPE1 cDNA |
| Lenti-EFS- BirA-GSQ-RNF4$^{\Delta SIM}$-P2A-BLAST | AMP | Lenti-EFS-GSQ-BirA-P2A-BLAST | RNF4$^{\Delta SIM}$ (BshT1-Asc1) | SIM-deficient version: 4xSIMs mutated (IELV, IVDL, PVVV and VVIV, all changed to AAAA) RNF4 Source: hTERT-RPE1 cDNA |
| Lenti-EFS- BirA-GSQ-MIB1-P2A-BLAST | AMP | Lenti-EFS-GSQ-BirA-P2A-BLAST | MIB1$^{WT}$ (BshT1-Asc1) | MIB1 Source: hTERT-RPE1 cDNA |
| Lenti-EFS- BirA-GSQ-MIB1$^{CA}$-P2A-BLAST | AMP | Lenti-EFS-GSQ-BirA-P2A-BLAST | MIB1$^{CA}$ (BshT1-Asc1) | Catalytically inactive: C963A mutation of 3$^{rd}$ RING domain. MIB1 Source: hTERT-RPE1 cDNA |
| Lenti-EFS- BirA-GSQ-MARCH5-P2A-BLAST | AMP | Lenti-EFS-GSQ-BirA-P2A-BLAST | MARCH5$^{WT}$ (BshT1-Asc1) | MARCH5 Source: hTERT-RPE1 cDNA |
| Lenti-EFS- BirA-GSQ-MARCH5$^{CA}$-P2A-BLAST | AMP | Lenti-EFS-GSQ-BirA-P2A-BLAST | MARCH5$^{CA}$ (BshT1-Asc1) | Catalytically inactive: C33A, C35A mutation of RING domain. MARCH5 Source: hTERT-RPE1 cDNA |
| Lenti-EFS- BirA-GSQ-RNF214-P2A-BLAST | AMP | Lenti-EFS-GSQ-BirA-P2A-BLAST | RNF214$^{WT}$ (BshT1-Asc1) | RNF214 Source: hTERT-RPE1 cDNA |

| **Plasmid Name** | **Res** | **Plasmid backbone** | **Plasmid insert** | **Sources/cloning notes** |
|---|---|---|---|---|
| Lenti-EFS- BirA-GSQ-RNF214$^{CA}$-P2A-BLAST | AMP | Lenti-EFS-GSQ-BirA-P2A-BLAST | RNF214$^{CA}$ (BshT1-Asc1) | Catalytically inactive: C675A/H677A/H680A/C683A mutation of RING domain. RNF214 Source: hTERT-RPE1 cDNA |
| Lenti-EFS-BirA-GSQ-NEDD4-P2A-BLAST | AMP | Lenti-EFS-BirA-GSQ-P2A-BLAST | NEDD4$^{WT}$ (EcoR1-Not1) | NEDD4 source: FLAG-NEDD4 plasmid, kind gift from S. Polo [10]. |
| Lenti-EFS-BirA-GSQ-NEDD4$^{CA}$-P2A-BLAST | AMP | Lenti-EFS-BirA-GSQ-P2A-BLAST | NEDD4$^{CA}$ (EcoR1-Not1) | Catalytically inactive, C867A mutation. NEDD4 source: FLAG-NEDD4 plasmid, kind gift from S. Polo [10]. |
| Lenti-EFS-BirA-GSQ-NEDD4$^{\Delta C2}$-P2A-BLAST | AMP | Lenti-EFS-BirA-GSQ-P2A-BLAST | NEDD4 $^{\Delta C2}$ (EcoR1-Not1) | Hyperactivated NEDD4, C2 domain deleted. NEDD4 source: FLAG-NEDD4 plasmid, kind gift from S. Polo [10]. |
| Lenti-EFS-BirA-GSQ-NEDD4$^{\Delta C2,CA}$-P2A-BLAST | AMP | Lenti-EFS-BirA-GSQ-P2A-BLAST | NEDD4 $^{\Delta C2, CA}$ (EcoR1-Not1) | Catalytically inactive (C867A) and C2 domain deleted. NEDD4 source: Lenti-EFS-BirA-GSQ-NEDD4$^{\Delta C2}$-P2A-BLAST. |
| Lenti-EFS-BirA-GSQ-NEDD4$^{3M}$-P2A-BLAST | AMP | Lenti-EFS-BirA-GSQ-P2A-BLAST | NEDD4$^{3M}$ (EcoR1-Not1) | Hyperactivated NEDD4, I36A, L37A and Y604A mutations. NEDD4 source: FLAG-NEDD4$^{3M}$ plasmid, kind gift from S. Polo [10]. |
| Lenti-EFS-BirA-GSQ-NEDD4$^{3M,CA}$-P2A-BLAST | AMP | Lenti-EFS-BirA-GSQ-P2A-BLAST | NEDD4$^{3M, CA}$ (EcoR1-Not1) | Catalytically inactive (C867A) and 3M mutant. NEDD4 source: Lenti-EFS-BirA-GSQ-NEDD4$^{3M}$-P2A-BLAST. |
| pcDNA3.1 (commercial) | AMP | pcDNA3.1 | empty | Invitrogen; transfection control |

**Supplementary Table 2: Oligonucleotides sequences and uses.** Related to Cloning section of *Methods*.

| Oligo name | Sequence (5'-3') | Used to construct following plasmids or intermediates |
|---|---|---|
| CMV_BIO_for | ACTCAGATCTCGAGCTCAAGCTTCGAATTC GCCACCATGGGTTTGAATGACATA | CMV-bioSUMO1$^{WT}$; CMV-bioSUMO2$^{WT}$; CMV-bioSUMO1nc; CMV-bioSUMO2nc |
| N1_SUMO1_wt_rev | ATGTGGTATGGCTGATTATGATCTAGAGTC GCGGCCGCTTAACCCCCCGTTTGTTCCTGAT A | CMV-bioSUMO1$^{WT}$ (used as template for TRIPZ versions) |
| N1_SUMO1_nocut_rev | GGTATGGCTGATTATGATCTAGAGTCGCGG CCGCTTAACCACCCGTAGGTTCCTGATAAA CTTCAATCACATCTTCTTC | CMV-bioSUMO1nc (used as template for TRIPZ versions) |
| N1_SUMO2_wt_rev | ATGTGGTATGGCTGATTATGATCTAGAGTC GCGGCCGCTTAACCTCCCGTCTGCTGTTGG AA | CMV-bioSUMO2$^{WT}$ (used as template for TRIPZ versions) |
| N1_SUMO2_nocut_rev | TGGTATGGCTGATTATGATCTAGAGTCGCG GCCGCTTAACCTCCCGTTGGCTGTTGGAAC ACATCAATTGTATCTTCAT | CMV-bioSUMO2nc (used as template for TRIPZ versions) |
| TRIPZ.EFS.for | CAGAGCTCGTTTAGTGAACCGTCAGATCGC TTGCCGCCAGAACACAGGACCGGT | To shuttle from Lenti-EFS vectors into TRIPZ |
| P2A.TRIPZ.rev | GCGCCAAAACCCGGCGCGGAGGCCACGCG TTCCGGCTTGTTTCAGCAGAGATCAGTTTGT TGCGCC | To shuttle from Lenti-EFS vectors into TRIPZ |
| TRIPZ.CMVorf.for | CAGAGCTCGTTTAGTGAACCGTCAGATCGC ACCGGTGCTGGTTTAGTGAACCGTCAGATC C | To amplify ORF-YFP (or other tags) inserts and stitch into Age1-Mlu1 digested TRIPZ |
| TRIPZ.marker.rev | CGGGAGGCGCCAAAACCCGGCGCGGAGGC CACGCGTGCTTTATTTGTGAAATTTGTGATG CTATTGCTTTATTTG | To amplify ORF-YFP-pA (or other tags, all have stops) inserts and stitch into Age1-Mlu1 digested TRIPZ |
| EFS.CMVorf.for | TTCGCAACGGGTTTGCCGCCAGAACACAGG ACCGGTGCTGGTTTAGTGAACCGTCAGATC C | To amplify ORF-YFP (or other tags) inserts and stitch into Age1-BamHI digested Lenti-EFS |
| P2A.CMVorf.rev | TTGTTTCAGCAGAGAGAAGTTTGTTGCGCC GGATCCCTTGTACAGCTCGTCCATGCCG | To amplify ORF-YFP (only; no stop) inserts and stitch into Age1-BamHI digested Lenti-EFS |
| EFS.seq.for | CGTATATAAGTGCAGTAGTCGCCGTGAACG TTC | To sequence 3' of EFS promoter |
| CMV-F | CGCAAATGGGCGGTAGGCGTG | To sequence 3' of CMV |
| Blast.seq.rev | GAGATGGGGATGCTGTTGATTGTAGCCG | To sequence 5' of blasticidin-resistance cassette |
| Puroseq.v2.rev | CCGGGGGACGTCGTCGCGGGTGG | To sequence 5' of puromycin-resistance cassette |
| GSQ.for | GGGCAAATTTCTTACGCAAGTAGAGGG | To sequence 3' of GSQ linker |
| TRIPZ.seqv2.for | GATGATTAATTGTCAACACGTGCTGCAGG | To sequence 3' of TRIPZ-tetO promoter |
| TRIPZ.seqv2.rev | CGTCTGACGTGGCAGCGCTC | To sequence 5' of TRIPZ Mlu1 site |
| Avitag.lowaff.for | GACATATTTGAAGCCCAGAAGATCGAGGGT GAGTTCGGCAGCGGCGAATTCATG | To modify Avitag (from WT to low affi version; WHE>GEF); works for bioSUMO1/2 |
| Avitag.lowaff.rev | CATGAATTCGCCGCTGCCGAACTCACCCTC GATCTTCTGGGCTTCAAATATGTC | To modify Avitag (from WT to low affi version; WHE>GEF); works for bioSUMO1/2 |
| Avitag.lowaffv2.for | GACATATTTGAAGCCCAGAAGATCGAGGGT GAGTTCGGATCCGGCTCCGGAATG | To modify Avitag (from WT to low affi version; WHE>GEF); works for bioUb |
| Avitag.lowaffv2.rev | CATTCCGGAGCCGGATCCGAACTCACCCTC GATCTTCTGGGCTTCAAATATGTC | To modify Avitag (from WT to low affi version; WHE>GEF); works for bioUb |

| Oligo name | Sequence (5'-3') | Used to construct following plasmids or intermediates |
|---|---|---|
| CMV_BIO_for | ACTCAGATCTCGAGCTCAAGCTTCGAATTC GCCACCATGGGTTTGAATGACATA | To amplify bioSUMO1, bioSUMO2 with wt or "no cut" C-term to stitch into CMV-R1-Not1 (Kan/Neo) |
| N1_SUMO1_wt_rev | ATGTGGTATGGCTGATTATGATCTAGAGTC GCGGCCGCTTAACCCCCCGTTTGTTCCTGAT A | To amplify bioSUMO1 with wt C-term to stitch into CMV-R1-Not1 (Kan/Neo) |
| N1_SUMO1_nocut_rev | ATGTGGTATGGCTGATTATGATCTAGAGTC GCGGCCGCTTAACCCCCCGTAGGTTCCTGA TA | To amplify bioSUMO1 with "no cut" C-term to stitch into CMV-R1-Not1 (Kan/Neo) |
| N1_SUMO2_wt_rev | ATGTGGTATGGCTGATTATGATCTAGAGTC GCGGCCGCTTAACCTCCCGTCTGCTGTTGG AA | To amplify bioSUMO2 with wt C-term to stitch into CMV-R1-Not1 (Kan/Neo) |
| N1_SUMO2_nocut_rev | TGGTATGGCTGATTATGATCTAGAGTCGCG GCCGCTTAACCTCCCGTTGGCTGTTGGAAC ACATCAATTGTATCTTCAT | To amplify bioSUMO2 with "no cut" C-term to stitch into CMV-R1-Not1 (Kan/Neo) |
| N1_UB_wt_rev | GGTATGGCTGATTATGATCTAGAGTCGCGG CCGCTTAACCACCTCTGAGACGGAGGACCA GGTGCAGGGT | To amplify bioUb with wt C-term to stitch into CMV-R1-Not1 (Kan/Neo) |
| N1_UB_nocut_rev | GGTATGGCTGATTATGATCTAGAGTCGCGG CCGCTTAACCACCTCTAGGACGGAGGACCA GGTGCAGGGT | To amplify bioUb with "no cut" C-term to stitch into CMV-R1-Not1 (Kan/Neo) |
| EFS.CEP120.for | CGGGTTTGCCGCCAGAACACAGGACCGGTG CCACCATGGTCTCCAAATCCGACCAATTGC TC | To amplify CEP120; to perform 3-way Gibson with BirAopt to generate L-EFS-CEP120-BirA-P2A-blast |
| BirA.CEP120.rev | TCAGGGGCACGGTGTTGTCCTTCATCGTCG ACTGATTACTGGCATTGCTTTTTGCCAAAAT CTC | To amplify CEP120; to perform 3-way Gibson with BirAopt to generate L-EFS-CEP120-BirA-P2A-blast |
| Sal1.BirAopt.for.v2 | CAGTCGACGATGAAGGACAACACCGTGCC CCTGA | To amplify BirAoptWT to make ORF-BirA fusions in EFS-P2A-blast (3-way Gibson) |
| P2A.BirA_orf.rev | TTGTTTCAGCAGAGAGAAGTTTGTTGCGCC GGATCCCTTCTCTGCGCTTCTCAGGGAGAT | To move X-BirA cassettes to EFS-2A-blast vector (3-way Gibson) |
| GSG.R1.PIAS1.for | CAAATTTCTTACGCAAGTAGAGGGGAATTC ATGGCGGACAGTGCGGAACTA | To amplify PIAS1 to stitch in Module2 (EcoR1-Not1); to make L-EFS-BirAopt-GSQ-PIAS1wt-P2A-blast |
| PIAS1.Not1.P2A.rev | GAAGTTTGTTGCGCCGGATCCGCGGCCGCC GTCCAATGAAATAATGTCTGGTATGATGCC | To amplify PIAS1 to stitch in Module2 (EcoR1-Not1); to make L-EFS-BirAopt-GSQ-PIAS1wt-P2A-blast |
| PIAS1.CA.for | CAATTCCGTGTCGGGCCCTTACAGCTTCTC ATCTACAATGTTTTGACG | To generate PIAS1 with mutated RING domain by 2-step PCR, Gibson. |
| PIAS1.CA.rev | CGTCAAAACATTGTAGATGAGAAGCTGTAA GGGCCCGACACGGAATTG | To generate PIAS1 with mutated RING domain by 2-step PCR, Gibson. |
| GSG.R1.PIAS4.for | CAAATTTCTTACGCAAGTAGAGGGGAATTC ATGGCGGACAGTGCGGAACTA | To amplify PIAS4 to stitch in Module2 (EcoR1-Not1); to make L-EFS-BirAopt-GSQ-PIAS1wt-P2A-blast |
| PIAS4.Not1.P2A.rev | GAAGTTTGTTGCGCCGGATCCGCGGCCGCC GCAGGCCGGCACCAGGCC | To amplify PIAS4 to stitch in Module2 (EcoR1-Not1); to make L-EFS-BirAopt-GSQ-PIAS1wt-P2A-blast |
| PIAS4.CA.for | CAAATTTCTTACGCAAGTAGAGGGGAATTC ATGGCGGCGGAGCTGGTG | To generate PIAS4 with mutated RING domain by 2-step PCR, Gibson. |
| PIAS4.CA.rev | GTCGAAGCACTGCAGGTGGGCGGCGGTCTC TGCCCGGCAGGG | To generate PIAS4 with mutated RING domain by 2-step PCR, Gibson. |
| GSQ.HsRNF4.for | CAAATTTCTTACGCAAGTAGAGGGGAATTC ATGAGTACAAGAAAGCGTCGTGGTGGA | To amplify RNF4 to stitch in Module2 (EcoR1-Not1); to make L-EFS-BirAopt-GSQ-RNF4wt-P2A-blast |
| P2A.HsRNF4.rev | GAGAGAAGTTTGTTGCGCCGGATCCGCGGC CGCCTATATAAATGGGGTGGTACCGTTTGT GG | To amplify RNF4 to stitch in Module2 (EcoR1-Not1); to make L-EFS-BirAopt-GSQ-RNF4wt-P2A-blast |
| HsRNF4.Nterm.dSIM.rev | GGCTCTAAAGATTCACAAGTAGCAGCCGCT GCTTCATCTCCAGCAGTTTCAGCTGCAGCG GCGGGTTCTGCTTCCAAGGA | To generate RNF4 without SIMs by 2-step PCR, Gibson. |

| Oligo name | Sequence (5'-3') | Used to construct following plasmids or intermediates |
|---|---|---|
| HsRNF4.Cterm.dSIM.for | GTGAATCTTTAGAGCCTGTGGCAGCTGCCGCAACTCACAATGACTCTGCTGCAGCAGCCGACGAAAGAAGAAGACCAAGG | To generate RNF4 without SIMs by 2-step PCR, Gibson. |
| HsRNF4.C159A.qc.for | CAGAATGCGGCCATGTCTTCGCTAGCCAGTGCCTCCGTGATTC | To generate RNF4 with mutated RING domain by 2-step PCR, Gibson. |
| HsRNF4.C159A.qc.rev | GAATCACGGAGGCACTGGCTAGCGAAGACATGGCCGCATTCTG | To generate RNF4 with mutated RING domain by 2-step PCR, Gibson. |
| GSQ_HsMIB1.for | CAAATTTCTTACGCAAGTAGAGGGGAATTCATGAGTAACTCCCGGAATAACCGGGTG | To amplify MIB1 to stitch in Module2 (EcoR1-Not1); to make L-EFS-BirAopt-GSQ-MIB1wt-P2A-blast |
| P2A_HsMIB1.rev | GAGAGAAGTTTGTTGCGCCGGATCCGCGGCCGCCATACAAAAGAATCCTTCGTTCAATAGCCTTGCG | To amplify MIB1 to stitch in Module2 (EcoR1-Not1); to make L-EFS-BirAopt-GSQ-MIB1wt-P2A-blast |
| HsMIB1.C963A.qc.for | CATTAAAGAGCAGACAATGGCCCCTGTGTGTCTAGATCG | To mutate 3rd RING domain of MIB1 by quickchange PCR method; catalytic-dead version |
| HsMIB1.C963A.qc.rev | CGATCTAGACACACAGGGGCCATTGTCTGCTCTTTAATG | To mutate 3rd RING domain of MIB1 by quickchange PCR method; catalytic-dead version |
| GSQ.HsPEX12.for | CAAATTTCTTACGCAAGTAGAGGGGAATTCATGGCTGAGCACGGGGCTCACTTCACA | To amplify PEX12 to stitch in Module2 (EcoR1-Not1); to make L-EFS-BirAopt-GSQ-PEX12wt-P2A-blast |
| P2A.PEX12.rev | GAGAGAAGTTTGTTGCGCCGGATCCGCGGCCGCCGTTCTCAGGGGAGTAGAGTTTAATCAGATG | To amplify PEX12 to stitch in Module2 (EcoR1-Not1); to make L-EFS-BirAopt-GSQ-PEX12wt-P2A-blast |
| PEX12.RING.CA.for | CCCCTCTTACCCAAAATGAAGACTGTGGCTCCACTGGCTCGTAAAACCCGGGTGAATGATACT | To mutate RING domain of PEX12 by quickchange PCR method; catalytic-dead version |
| PEX12.RING.CA.rev | AGTATCATTCACCCGGGTTTTACGAGCCAGTGGAGCCACAGTCTTCATTTTGGGTAAGAGGGG | To mutate RING domain of PEX12 by quickchange PCR method; catalytic-dead version |
| GSQ.HsMARCHF5.for | CAAATTTCTTACGCAAGTAGAGGGGAATTCATGCCGGACCAAGCCCTACAGCAG | To amplify MARCHF5 to stitch in Module2 (EcoR1-Not1); to make L-EFS-BirAopt-GSQ-MARCHF5wt-P2A-blast |
| P2A. HsMARCHF5.rev | GAGAGAAGTTTGTTGCGCCGGATCCGCGGCCGCCTGCTTCTTCTTGTTCTGGATAATTCAGAAT | To amplify PEX12 to stitch in Module2 (EcoR1-Not1); to make L-EFS-BirAopt-GSQ-MARCHF5wt -P2A-blast |
| HsMARCHF5.RING.CA.for | ACAGCTGAATGGGTGAGACCAGCCAGGGCTAGAGGATCTACAAAATGGGTT | To mutate RING domain of MARCHF by quickchange PCR method; catalytic-dead version |
| HsMARCHF5.RING.CA.rev | AACCCATTTTGTAGATCCTCTAGCCCTGGCTGGTCTCACCCATTCAGCTGT | To mutate RING domain of MARCHF5 by quickchange PCR method; catalytic-dead version |
| GSQ.HsRNF214.for | CAAATTTCTTACGCAAGTAGAGGGGAATTCATGGCAGCGTCTGAGGTTGCTGGTGTT | To amplify PEX12 to stitch in Module2 (EcoR1-Not1); to make L-EFS-BirAopt-GSQ-RNF214wt-P2A-blast |
| P2A.HsRNF214.rev | GAGAGAAGTTTGTTGCGCCGGATCCGCGGCCGCCTTTAAGAGTTGGACAAAAGGGACAAGTGTC | To amplify PEX12 to stitch in Module2 (EcoR1-Not1); to make L-EFS-BirAopt-GSQ- RNF214wt-P2A-blast |
| HsRNF214.RING.CA.for | GCGGCTACCGCTGTATTGGCCAAGGAGGCTATCAAATTCTGGGCCCAGACC | To mutate RING domain of RNF214 by 2-step overlap PCR method; catalytic-dead version |
| HsRNF214.RING.CA.rev | GATAGCCTCCTTGGCCAATACAGCGGTAGCCGCCATTGGATGCAGCTCACT | To mutate RING domain of RNF214 by 2-step overlap PCR method; catalytic-dead version |
| GSQ.NEDD4.for | CAAATTTCTTACGCAAGTAGAGGGGAATTCATGGCAACTTGCGCGGTGGAGGTG | To amplify NEDD4 to stitch in Module2 (EcoR1-Not1); to make L-EFS-BirAopt-GSQ-NEDD4wt-P2A-blast (E. Maspero clone as template; includes N-term C2 domain) |
| P2A.NEDD4.rev | GAGAGAAGTTTGTTGCGCCGGATCCGCGGCCGCCATCAACTCCATCAAAGCCCTGGGTGTT | To amplify NEDD4 to stitch in Module2 (EcoR1-Not1); to make L-EFS-BirAopt-GSQ-NEDD4wt-P2A-blast (E. Maspero clone as template; includes N-term C2 domain) |
| NEDD4.C1286A.for | AAGCTGCCAAGAGCTCATACCGCTTTTAATCGCCTGGACTTGCCA | To mutate HECT domain thioester transfer Cys>Ala |

| Oligo name | Sequence (5'-3') | Used to construct following plasmids or intermediates |
|---|---|---|
| NEDD4.C1286A.rev | TGGCAAGTCCAGGCGATTAAAAGCGGTATG AGCTCTTGGCAGCTT | To mutate HECT domain thioester transfer Cys>Ala |
| NEDD4.stop.for | CCCAGGGCTTTGATGGAGTTGATTAGTAAG GCGGCCGCGGATCCGGC | To introduce STOP codon at C-terminus of NEDD4 to avoid leaving extended P2A remnant |
| NEDD4.stop.rev | GCCGGATCCGCGGCCGCCTTACTAATCAAC TCCATCAAAGCCCTGGG | To introduce STOP codon at C-terminus of NEDD4 to avoid leaving extended P2A remnant |
| GSQ.NEDD4dC2.for | CGGGTTTGCCGCCAGAACACAGGACCGGTG CCACCATGACTTATTTACCTAAAACCAGTG GCTCA | To amplify NEDD4 without C domain (as in NM_001329212; E. Maspero clone as template; lacks N-term C2 domain) |
| NEDD4.700.for | GCAACATTCAACTGCAAGCACAACG | For full sequencing of NEDD4 |
| NEDD4.1400.for | TACTTCCAATGATCTAGGGCCTTTACC | For full sequencing of NEDD4 |
| NEDD4.2100.for | CATAGATGAAGAACTTTTTGGACAGACAC | For full sequencing of NEDD4 |
